# The *Synchytrium endobioticum* AvrSen1 triggers a Hypersensitive Response in Sen1 potatoes while natural variants evade detection

**DOI:** 10.1101/646984

**Authors:** Bart T.L.H. van de Vossenberg, Charlotte Prodhomme, Gert van Arkel, Marga P.E. van Gent-Pelzer, Marjan Bergervoet, Balázs Brankovics, Jarosław Przetakiewicz, Richard G.F Visser, Theo A.J. van der Lee, Jack H. Vossen

**Author notes:** Corresponding author (JV).

## Abstract

*Synchytrium endobioticum* is an obligate biotrophic fungus of the phylum Chytridiomycota. It causes potato wart disease, has a world-wide quarantine status and is included on the HHS and USDA Select Agent list. *S. endobioticum* isolates are grouped in pathotypes based on their ability to evade host-resistance in a set of differential potato varieties. So far, thirty-nine pathotypes are reported. A single dominant gene (*Sen1*) governs pathotype 1 resistance and we anticipated that the underlying molecular model would involve a pathogen effector (AvrSen1) that is recognized by the host. The *S. endobioticum* specific secretome of fourteen isolates representing six different pathotypes was screened for effectors specifically present in pathotype 1(D1) isolates but absent in others. We identified a single *AvrSen1* candidate. Expression of this candidate in potato *Sen1* plants showed a specific hypersensitive response, which co-segregated with the *Sen1* resistance in potato populations. No HR was obtained with truncated genes found in pathotypes that evaded recognition by *Sen1*. These findings established that our candidate gene was indeed *Avrsen1.* The *S. endobioticum AvrSen1* is a single copy gene and encodes a 376 amino acid protein without predicted function or functional domains, and is the first effector gene identified in Chytridiomycota, an extremely diverse yet underrepresented basal lineage of fungi.

**Author Summary:** Plant pathogens can have a great social and economic impact, and are a continuous threat to food security. A clear example is *Synchytrium endobioticum*, the fungus causing potato wart disease. The impact of the pathogen, lack of effective chemical control agents and the longevity of resting spores produced by the pathogen led to a world-wide quarantine status for *S. endobioticum*. Strict phytosanitary measures and the use of resistance potato varieties are currently the only way to prevent the spread of the disease. The emergence of new pathotypes that overcome resistance urged to study the underlying molecular mechanisms of *S. endobioticum* recognition by the plant. Here we describe the identification of the first effector (AvrSen1) of *S. endobioticum* that is recognized by the *Sen1* resistance gene product. Also, we report the loss of *AvrSen1* in other pathotypes thus avoiding recognition by the plant and triggering immune responses. AvrSen1 represents the first effector to be identified in the basal fungal lineage Chytridiomycota. The discovery of AvrSen1 provides an important tool to manage potato wart disease. Moreover, knowledge about Chytridiomycota effectors will shed light on other (pathogenic) interactions and the co-evolution of Chytridiomycota species with their hosts.

## Introduction

Potato wart is a severe disease of cultivated potatoes, caused by the soil borne obligate biotrophic fungus *Synchytrium endobioticum* (Schilb.) Percival. Hypertrophic growth of the infected tissue resulting in wart-like malformations that destroy the economic value of the potato tubers, characterize the disease (1). Resting spores that are formed in the warted potato tissues can remain viable and infectious in soils for decades (2). No chemical control agents are available to eradicate the pathogen from contaminated soils (3), and disease management relies on preventing its introduction and spread through the deployment of resistant potato varieties. Because of its impact on potato cultivation, *S. endobioticum* has a quarantine status in most countries with potato production and is included on the HHS and USDA Select Agent list (4). The pathogen has been reported in potato-growing countries in Asia, Africa, Europe, North-America, South-America, and Oceania (5).

Isolates of the pathogen are grouped into pathotypes based on their interaction with a differential set of resistant potato varieties (6). For decades after the first description of the pathogen by Schilberszky (7), only a single pathotype was recognized, nowadays referred to as pathotype 1(D1). Emergence of a new pathotype, now known as 2(G1), was recognized when wart formation was discovered on formerly resistant potato cultivars in 1941 (8). Currently, 39 pathotypes have been described, of which pathotypes 1(D1), 2(G1), 6(O1) and 18(T1) are most widespread in western Europe and considered to be of main importance (9). Comparative analysis of thirty *S. endobioticum* isolates using their mitochondrial genome sequences, showed that the pathogen was introduced multiple times in Europe and successively several pathotypes emerged in parallel. Interestingly, single isolates we found to represent populations of distinct genotypes (10).

*S. endobioticum* belongs to the Chytridiomycota, a basal lineage of fungi that are evolutionary diverse and arose in the Mesoproterozoic Era about 1,000 to 1,600 million years ago (11). Despite being ubiquitous in nature, only a few Chytridiomycota are studied at the molecular or genomic level. There are approximately 1,500 formally described chytrid species and the genus *Synchytrium* alone contains over 200 described species, most of which are obligate biotrophic plant pathogens. *S. endobioticum* is one of the best studied Chytridiomycota, but thus far studies on *S. endobioticum* focused on its life cycle, epidemiology, pest management, and molecular tools for detection (reviewed in (12, 13)). Little is known about the molecular mechanisms underlying the obligate biotrophic or pathogenic lifestyle of this pathogen. As other Chytridiomycota, *S. endobioticum* does not form hyphae or mycelia but produces summer and resting sporangia that contain motile zoospores (14, 15). Zoospores encyst on the potato host cell and the content of the spore penetrates the host cell leaving the empty cyst wall outside the host. After penetration, the fungal thallus is separated from the point of infection and migrates to the host nucleus. The intracellular lifecycle is completed by forming summer sporangia which give rise to new zoospores that either re-infect the host or conjugate to produce biflagellate zygotes that give rise to resting spores after host penetration (1, 16, 17). Even in incompatible interactions, zoospores have been reported to penetrate host cells after which an immune response is triggered resulting in a localized cell death (18).

Plant - pathogen interactions have evolved over millions of years, generating a broad range of diversity on both sides of the interaction The molecular mechanisms involved in plant-fungi interactions have been reviewed by various authors (19-22). Weapons in this arms race are pathogen effector proteins and plant resistance (*R*) genes. Pathogen genes coding for effectors that are recognized by a plant *R* gene and trigger effector-triggered immunity (ETI) are called avirulence (*Avr*) genes. In agricultural systems the arms race model (23), in which both the pathogen and the host develop in continuous cycles causing temporary fixation of new effector and *R* gene alleles, has been suggested to be the main driving force in pathogen effector and plant defense evolution (24).

Potato is host to many pathogens from diverse taxonomical groups, such as oomycetes (e.g. *Phytophthora infestans*), bacteria (e.g. *Ralstonia* species), nematodes (e.g. *Globodera species*), but also viruses (e.g. PVY). Resistance in most pathosystems is governed by *R* gene recognition of specific effector molecules. The best elaborated examples are the gene-for gene (25) interactions in the *P. infestans* pathosystem (26). In potato, several quantitative resistance loci (QRLs) for *S. endobioticum* resistance have been identified, and two of these give resistance to pathotype 1(D1) isolates only: *Sen1* (27) and *Sen1-4* (28) which reside on chromosomes 11 and 4 respectively. Recently, *Sen2* and *Sen3* were described which give broad resistance to multiple pathotypes, including 1(D1) for *Sen2* (29, 30).

The recently annotated genomes of the pathotype 1(D1) isolate MB42 (QEAN00000000 v.1) and pathotype 6(O1) isolate LEV6574 (QEAM00000000 v.1) (31) open up the possibility to identify *Avr* gene candidates using a comparative genomic approach. Availability of *S. endobioticum Avr* genes will greatly advance our understanding and management of this challenging obligate biotrophic soilborne pathogenic fungus. Comparative studies of genome sequences from *S. endobioticum* isolates most frequently found in Europe and Canada, revealed an *AvrSen1* gene candidate which was present in single copy in pathotype 1(D1) genomes. The gene showed different variants in pathotypes that are not recognized by *Sen1*. This gene represents the first avirulence gene reported from a pathogen in the fungal phylum of Chytridiomycota. We discuss the potential applications of the applied comparative genomic strategy and the identified *AvrSen1* gene for potato wart disease resistance and management.

## Results

### *Screening for AvrSen1* candidates

Out of sequence data generated by Van de Vossenberg *et al*. (10), fourteen isolates were selected because of their >10 x median sequence coverage and known pathotype identity (Fig. S1). The *S. endobioticum* pathotype 1(D1) isolate MB42 genome contains 8,031 protein coding genes of which 477 (5.9 %) were regarded as the MB42 secretome on account of the presence of a signal peptide and absence of transmembrane domains or GPI anchors. Almost two-thirds of the secretome (n =304, 64 %) consists of species specific genes (Table S3).

To determine polymorphisms at the encoded protein sequence level (loss of function) and the DNA level (gene loss), reads of the 14 isolates were mapped to all nuclear genome scaffolds of pathotype 1(D1) isolate MB42. For the different isolates, dN substitutions were observed in 171 to 1,126 gene models at a minimal frequency of 70% (Table S4). Pathotype 8(F1), 18(T1) and 38(N1) isolates had significantly higher numbers of genes with dN substitutions (general ANOVA *p*<0.001). From the structural absence analysis of the different isolates, 41 to 206 gene models had less than 90% coverage relative to the MB42 reference genome and were predicted to be structurally absent.

Five gene models showed the hypothesized *AvrSen1* pattern, i.e. present in pathotype 1(D1) isolates and absent in the higher pathotypes. Four of these genes were species specific, and only one belonged to the secretome (Fig. 1). Manual verification of gene prediction, functional annotations, and weighing of the significance of polymorphisms (i.e. leading to conservative or non-conservative non-synonymous substitutions), only a single gene remained as a *AvrSen1* candidate: SeMB42_g04087. Opposed to the other genes with the hypothesized *AvrSen1* pattern, all polymorphisms in SeMB42_g04087 led to the introduction of stop codons resulting in truncated gene models.

**Figure 1.**
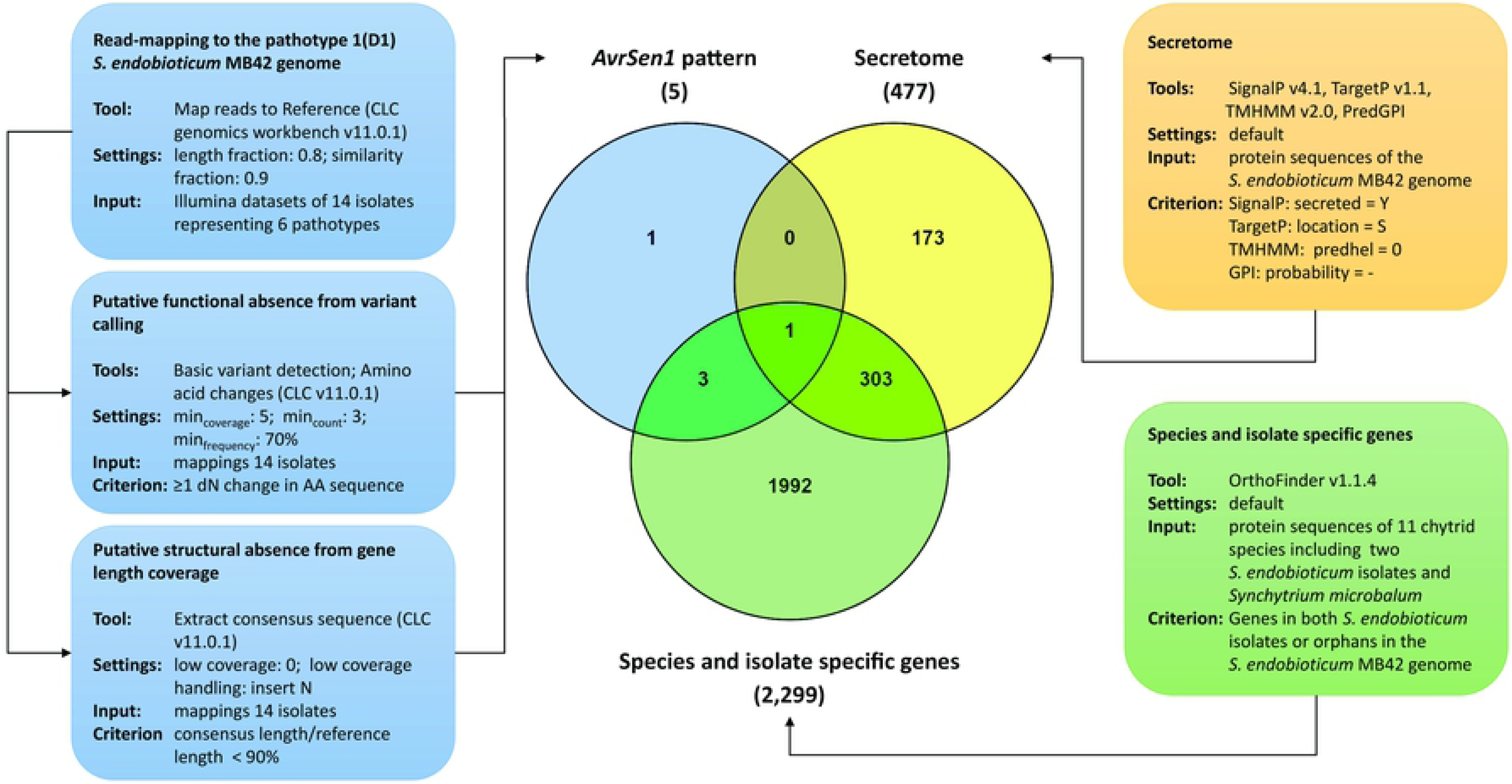
Converging approaches for the identification of *AvrSen1* candidates. The pipeline contains three main elements indicated in blue (putative presence and absence of genes), yellow (determination of the secretome) and green (species and isolate specific genes). Bioinformatic tools, their settings, input and output criteria are shown. Intersections of these three elements resulted in the identification of a single candidate *AvrSen1* gene from a total of 8,031 genes.

Gene SeMB42_g04087 is unique to *S. endobioticum* genomes, even in the closely related species *S. microbalum* no orthologs were found. The gene is 1,360 nucleotides long, consists of four exons and three introns, and encodes a 376 amino acid protein containing a signal peptide with cleavage site at amino acid position 30. No predicted functions, transmembrane domains, or other functional annotations such as nuclear localization signals, chloroplast targeting peptides or mitochondrial targeting peptides were identified. The protein contains two cysteine residues, and compared to the entire *S. endobioticum* MB42 proteome it has a relative high percentage of tyrosine residues (MB42 proteome: 2.8%; SeMB42_g04087: 5.1%), while other amino acid residues show more average frequencies.

### *AvrSen1* and its variants

The candidate *AvrSen1* gene was found to be present in the two pathotype 1(D1) isolates, and structurally absent in one of two pathotype 2(G1) isolates (i.e. MB08) and in both pathotype 18(T1) isolates (Fig. 2). In the other isolates two forms of functional absence were observed: a G insertion at position 769 of the coding sequence (CDS) causing a frameshift, thereby introducing a stop codon at position 256 of the amino acid sequence (avrSen1:Asp^256^>stop^256^); and a C>T substitution at position 916 of the CDS causing the introduction of a stop codon at position 306 of the amino acid sequence (avrSen1:Gln^306^>stop^306^). avrSen1:Asp^256^>stop^256^and avrSen1:Gln^306^>stop^306^ are referred to as variant 1 and variant 2 respectively (Fig. 2). Variant 1 was detected in pathotype 2(G1) isolate SE4, 6(O1) isolates SE5 and SE6, 8(F1) isolate DEN01, and 38(N1) isolate MB56. Variant 2 was exclusively found in the Canadian pathotype 6(O1) isolates LEV6574, LEV6602, LEV6687, and LEV6748. Also, two forms of structural absence were identified: i.e. a deletion ranging from position 29,142 to 30,137 (SeMB42scf158 Δ29,142-30,137) which was found in the two pathotype 18(T1) isolates (MB17 and SE7), and a deletion found only in one pathotype 2(G1) isolate (MB08) (SeMB42scf158Δ28,298-30,137). These two forms of structural absence are referred to as variants 3 and 4 respectively.

**Figure 2.**
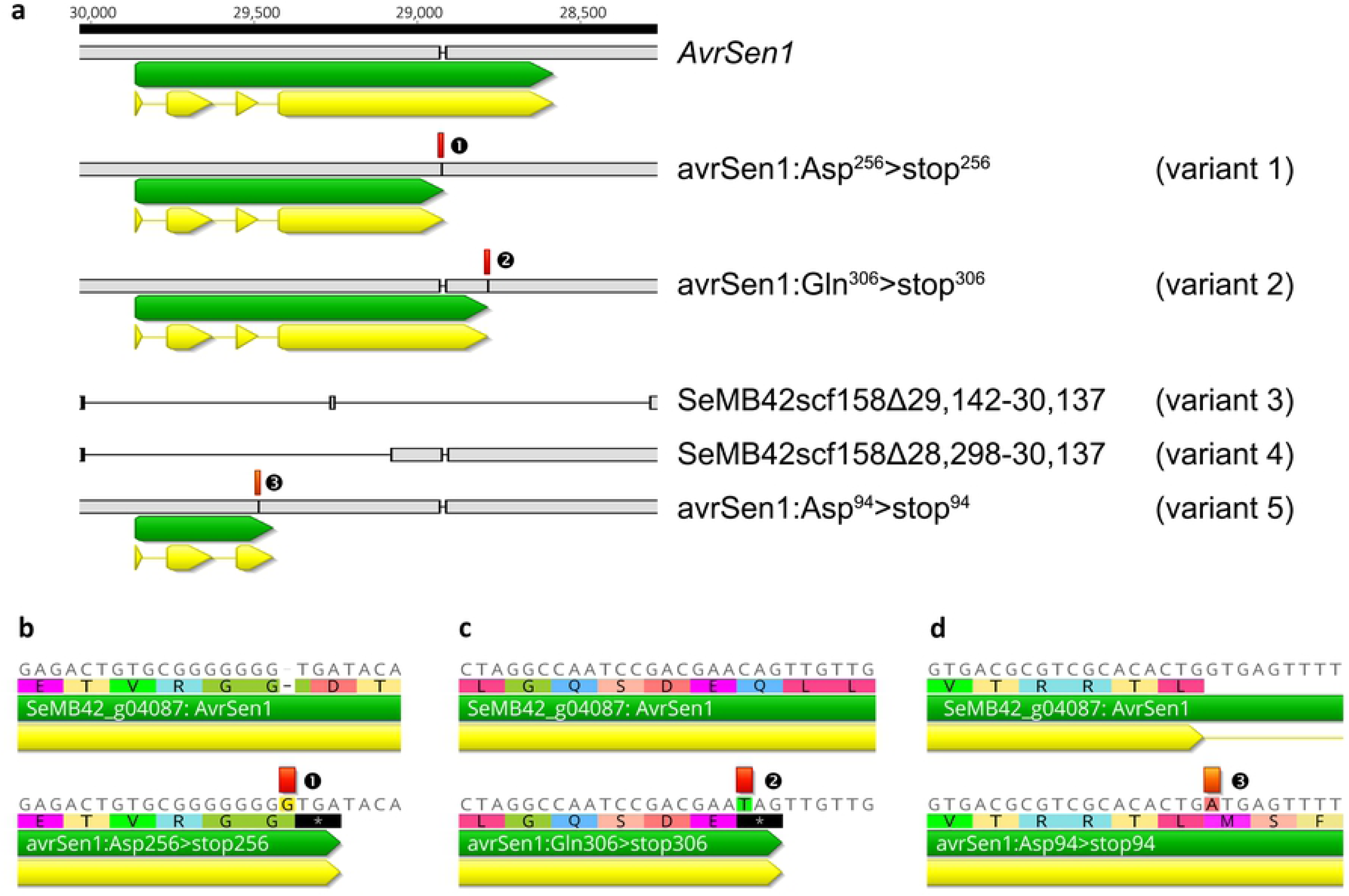
*AvrSen1* gene and its variants. **A** Sequences identical to the MB42 *AvrSen1* gene are presented in light grey, and variants to the *AvrSen1* gene are annotated in red (❶, ❷ and ❸ for the respective truncated variants). Gene annotation and coding sequence (CDS) annotation are in green and yellow respectively. **B** Detail of the G insertion in the genomic sequence of variant 1 isolates relative to isolate MB42 at position 769 of the CDS causing a frameshift and introducing a stop codon. **C** Detail of the C>T substitution in the genomic sequence of variant 2 isolates relative to isolate MB42 at position 916 of the CDS introducing a stop codon. **D** Detail of the G>A substitution in the genomic sequence of the variant 5 isolate on the first base of the third intron as present in isolate MB42 which results in a loss of the splice site. The numbering above the alignment indicates the original position on SeMB42_scf158 as it is represented in reverse complement orientation.

### Specific recognition of AvrSen1 in potato clones carrying the *Sen1* locus

Eight progeny plants were selected from the Aventra x Desiree and Aventra x Kuras crosses. Together with the parents of these populations, seven genotypes possessed the *Sen1* markers and were resistant to *S. endobioticum* pathotype 1(D1) whereas the four remaining genotypes lacked the *Sen1* markers and were susceptible to pathotype 1(D1). In total 153 agroinfiltrations were performed with the *AvrSen1* construct without signal peptide (AvrSen1ΔSP) in four different experiments, 45 with the *AvrSen1* construct with signal peptide (AvrSen1+SP), and 24 for each of the truncated variants without signal peptide (Table S5). When plants possessing *Sen1* were agroinfiltrated with the AvrSen1ΔSP construct, HR-like cell death was visible in 95 % of all infiltrated sites, this in contrast to agroinfiltration of AvrSen1ΔSP in plants lacking *Sen1*, where 98 % produced no visible reaction. This clearly showed that AvrSen1 was recognized in a Sen1 dependent way. Interestingly, the AvrSen1+SP construct did not produce an HR in *Sen1* plants suggesting cytoplasmic recognition of the gene product. Also, both truncated variants avrSen1:Asp^256^>stop^256^ and avrSen1:Gln^306^>stop^306^ were not recognized by *Sen1* plants and produced no visible reaction (Fig. 3).

**Figure 3.**
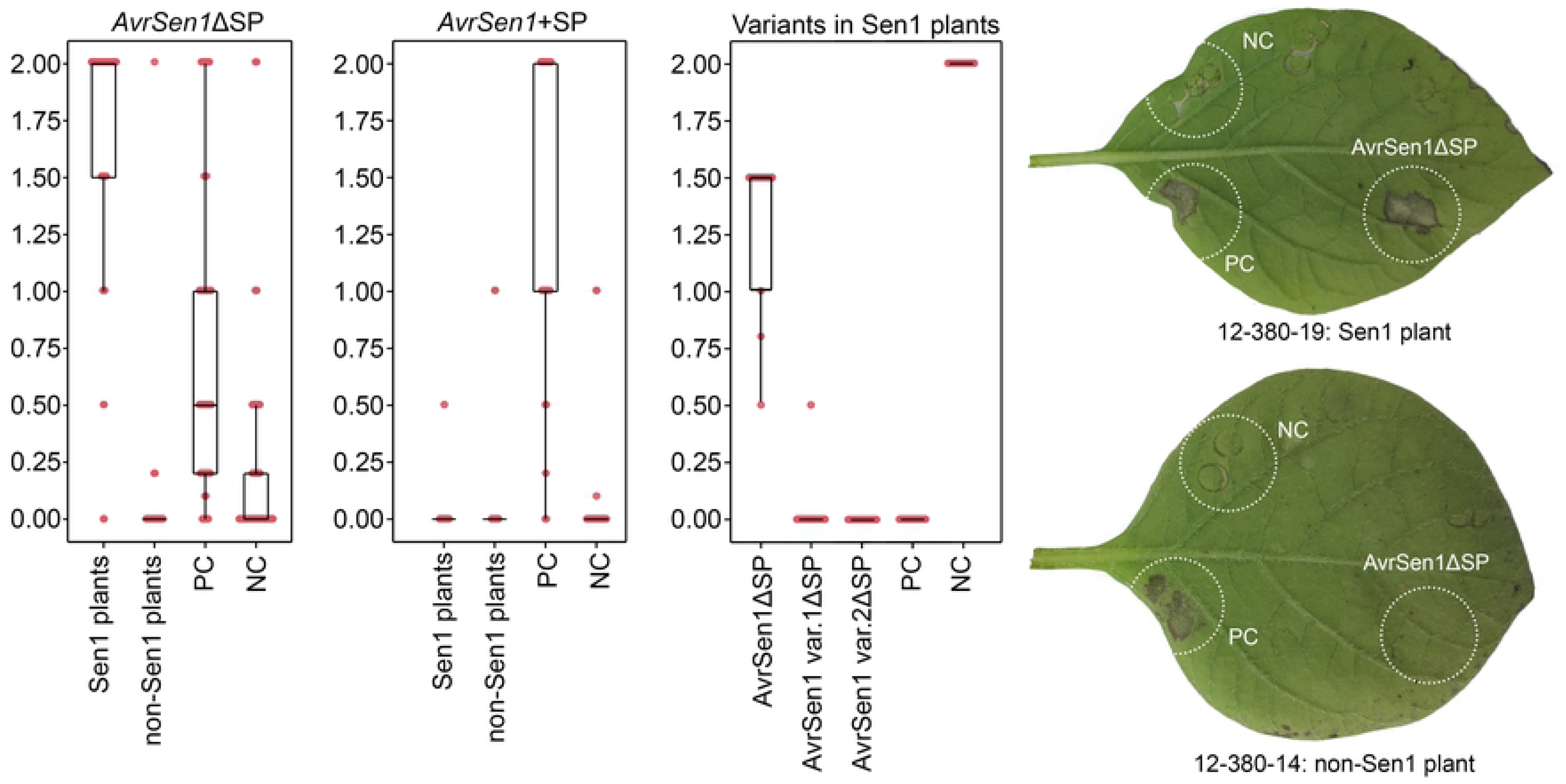
Agroinfiltration of *AvrSen1* variants in potato plants. Boxplots of Agroinfiltration scores obtained with constructs *AvrSen1*ΔSP (left panel) in which *Avr8/R8* co-infiltration served as positive control, *AvrSen1*+SP (center panel) in which *Avr8/R8* coinfiltration served as positive control, and the truncated *AvrSen1* variants 1 and 2 ΔSP (right panel) in which *AvrSen1*ΔSP and *Avr8*/*R8* co-infiltration served as positive controls. The negative control consisted of GUS or *R8*. Individual scores are represented as dots. Agroinfiltration results obtained in two progeny plants of the Aventra x Kuras population represent typical reactions observed for *AvrSen1*ΔSP in *Sen1* containing plants (top leaf) and plants without *Sen1* (bottom leaf). PC represents the positive control which consisted of *Avr8*/*R8* co-infiltration, and NC represents the negative control (GUS).

### Presence of *AvrSen1* and its variants in *S. endobioticum* isolates

The presence of the *AvrSen1* gene or its variants could be determined in seventeen of thirty *S. endobioticum* isolates sequenced by Van de Vossenberg *et al*. (10) by mapping of Illumina sequence reads to the MB42 genome. In addition, PacBio SMRT CCS reads generated from the *AvrSen1* amplicon allowed the detection of *AvrSen1* gene or its variants for three additional isolates. Totally, for 20 isolates the major *AvrSen1* variant was determined and superimposed to the mitochondrial haplotype network that was previously generated (Fig. 4). Based on variation of the mitochondrial genome, *S. endobioticum* isolates clustered in four major groups (haplogroups), representing separate introductions from which pathotypes 2(G1) and 6(O1) emerged multiple times independently (10).

**Figure 4.**
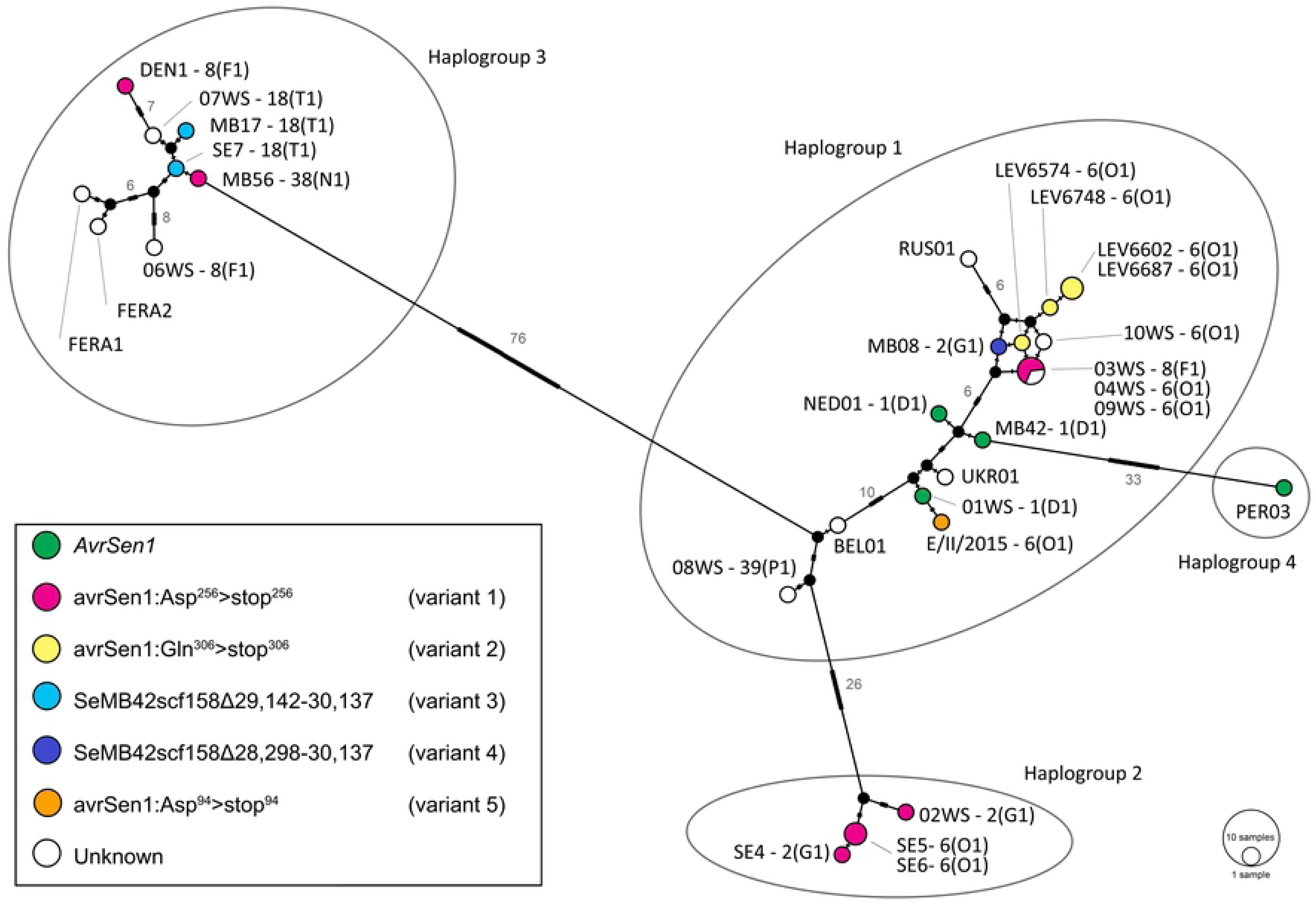
Distribution of *AvrSen1* and its variants among *S. endobioticum* isolates visualized in the mitochondrial haplotype network from Van de Vossenberg *et al*. (10). Colors represent the presence of *AvrSen1* or its variants for a given isolate as identified from read-mapping to the MB42 genome or through PacBio amplicon sequencing. Pathotype identities are shown when available. Black nodes represent hypothetical ancestors, and marks on the branches indicate the number of mutations, which are shown as numerical values on branches with > 5 mutations.

The *AvrSen1* gene was identified in both pathotype 1(D1) isolates from the Netherlands (MB42 and NED1), the German pathotype 1(D1) isolate 01WS, and in the Peruvian isolate without known pathotype identity. Variant 1 of *AvrSen1* was found most frequently among the remaining isolates, and was identified in haplogroups 1, 2 and 3. Interestingly, pathotype 6(O1) samples from Canada possess a different *AvrSen1* variant compared to the European pathotype 6(O1) isolates, i.e. variant 2 versus variant 1 respectively. Also, pathotype 2(G1) isolate MB08 from haplogroup 1 possesses a different variant compared to the pathotype 2(G1) isolates from haplogroup 2, i.e. variant 4 compared to variant 1 respectively.

### Co-occurrence of *AvrSen1* and its variants within *S. endobioticum* isolates

Read mappings generated of several isolates for the identification of the *AvrSen1* candidates showed that the *AvrSen1* locus was polymorphic in some isolates, and low percentages of the *AvrSen1* wild-type were present in these isolates. To further quantify the presence of *AvrSen1* and its variants within isolates, the gene was PCR amplified from genomic DNA (Fig. 5A, B) and subjected to PacBio SMRT sequencing which produced 8,793 to 32,336 CCS for the selected isolates (Table S1). PacBio CCS reads were generated from a median number of 36 or 37 passages for all isolates included (Fig. S3). In pathotype 1(D1) isolate MB42, all reads represented the *AvrSen1* wild-type (Fig. 5C). PacBio SMRT sequencing confirmed the presence of variant 1 as the dominant haplotype in isolates SE4, SE5, SE6, and MB56 that were also included in the screening to identify *AvrSen1* candidates. In these isolates, representing three higher pathotypes, the G insertion at position 769 of the coding sequence was observed in 93% of all reads. Interestingly, the *AvrSen1* haplotype was also found in these samples with 7% of the reads lacking the insertion that leads to the truncation of the gene. Pathotype 2(G1) isolate 02WS, which was not included in the screening for *AvrSen1* candidates because a lack of read coverage, showed presence of the *AvrSen1* variant 1 insertion in similar percentages (92%) as the other isolates carrying the *AvrSen1* variant 1 sequenced with PacBio.

**Figure 5.**
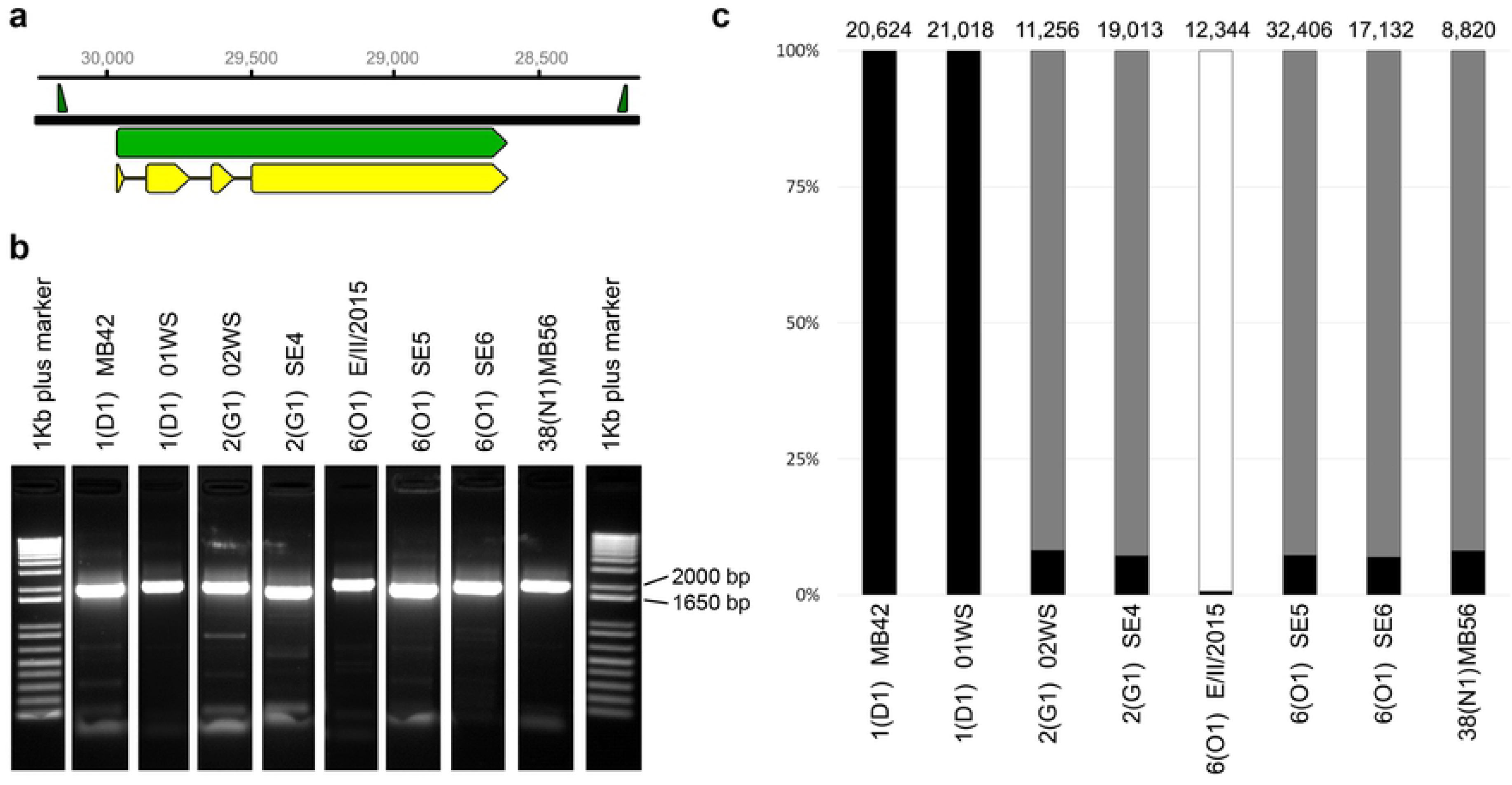
*AvrSen1* amplicon sequencing with PacBio SMRT. **A** Primer sites flanking the *AvrSen1* gene resulting in a 1,983 nt amplicon. Primers are annotated as green triangles, the *AvrSen1* gene sequence is annotated green, and the CDS is annotated in yellow. Numbers indicate the position of the gene on SeMB42_scf158 which is presented in reverse complement orientation to present the gene in 5’ to 3’ direction. **B** Amplicons obtained from selected isolates representing multiple pathotypes and mitochondrial haplogroups. The 1 kb-plus size marker was used for amplicon size estimation. **C** Proportion of PacBio CCS reads representing *AvrSen1* (black), *avrSen1* variant 1 (grey) and *avrSen1* variant 5 (white). The number of PacBio CCS reads generated per sample are shown above the bars.

In addition to the reads mapping to the MB42 genomic scaffold containing *AvrSen1*, low numbers (1 - 57) of small (95 - 640 nt) CCS reads were obtained for the tested isolates. In many cases these small CCS reads could be identified as non-specific amplification of bacteria or potato. However, a particular 466 to 470 bp amplicon representing a deletion variant was found to be present in six of the analyzed isolates in 2 to 5 copies per isolate (on average 1.6 in 10,000 reads) (Fig. S4). The short sequences resemble the *AvrSen1* variant 3 deletion, and were found in both 1(D1) isolates, both 2(G1) isolates, and the 6(O1) isolates SE5 and SE6. The short sequence was not identified in 38(N1) isolate MB56 nor in 6(O1) isolate E/II/2015.

### Disruptive selection

After two multiplication rounds of pathotype 1(D1) isolate 01WS on the semi-susceptible variety Erika, a pathotype 6(O1) phenotype was obtained for the resulting isolate E/II/2015 (10). Both isolates 01WS and E/II/2015 were PacBio sequenced to determine if a shift in the genetic population had resulted in the loss of the *AvrSen1* gene.

Almost all (>99.9%) of the 20.986 PacBio CCS reads obtained for pathotype 1(D1) isolate 01WS represented the *AvrSen1* haplotype. The *AvrSen1* haplotype in E/II/2015 was almost completely lost with only 0.7% of all 12,322 reads producing the wild-type sequence. The remaining 99.3% of the reads showed a G>A substitution on position 29,183 of scaffold SeMB42_scf0158 (Fig. S5). In the original 01WS isolate, seven CCS reads had the G>A substitution (0.0003%) observed in E/II/2015. This substitution is positioned on the first base of the third intron and affects the GU dinucleotide of the third exon-intron boundary in the pre-mRNA. The conserved GU dinucleotide is required for correct splicing of the pre-mRNA and the G>A substitution in the pre-mRNA results in a loss of the splice site. As a consequence, a stop codon is introduced on position 94 of the amino acid sequence resulting in a third truncated variant (avrSen1:Asp^94^>stop^94^); i.e. variant 5 (Fig. 2).

## Discussion

Close to forty different *S. endobioticum* pathotypes have been reported based on bioassays using a set of differentially resistant potato varieties. We hypothesized that the phenotypic differences between pathotypes are the result of different (combinations of) *R* genes present in potato varieties and presence or absence of their cognate *Avr* genes in *S. endobioticum* isolates. At present, several QRLs providing resistance to *S. endobioticum* pathotypes have been described of which two provide resistance specifically to pathotype 1(D1) isolates: i.e. *Sen1* and *Sen1-4*. To identify the cognate *Avr* for the potato *Sen1*, which we refer to as *AvrSen1*, we exploited the recently assembled and annotated pathotype 1(D1) genome of isolate MB42 in combination with Illumina datasets generated for 30 isolates representing pathotypes that are of current importance in Europe or Canada.

### *AvrSen1* identification strategy

As potato varieties possessing the *Sen1* locus are resistant to pathotype 1(D1) isolates but are susceptible to the “higher” pathotypes, we hypothesized that pathotype 1(D1) isolates possess the *AvrSen1* gene while higher pathotypes lost it on a functional or structural level. For the comparative genomic approach, two scenarios of loss of the *AvrSen1* gene were considered: mutations in the gene sequence leading to a loss of recognition due to the change in amino acid sequence of the AvrSen1 protein, and (partial) deletion of the gene from the genome in higher pathotypes. We did not include criteria such as molecular weight, size and cysteine-richness which have been attributed to be effector signatures (32). Indeed, effectors frequently are reported to be cysteine-rich and relatively small, but large effector proteins have also been reported (20). Also selection for other effector signatures based on specific motifs (e.g. the RxLR motif in oomycetes (33) or Crinkler motifs which were also found identified in the amphibian decline chytrid *B. dendrobatidis* (34)), were not pursued in our approach.

Only five candidates were obtained from the presence/absence analysis, of which only one was predicted to be secreted. Manual verification showed that the gene models of the other candidates were correct and that these indeed did not belong to the secretome. Otherwise, manual verification was found to be essential to assess the reliability of read-mapping and the resulting variant calling. Also, the significance of the dN changes were manually assessed with conservative dN changes, resulting in an amino acids with the same characteristics (e.g. Alanine to Valine), being regarded as insignificant as their influence to the tertiary structure of the protein was believed to be minimal. Notably, all single nucleotide polymorphisms observed in the *AvrSen1* gene of non-pathotype 1(D1) isolates led to the introduction of a stop codon resulting in a truncated gene model.

Pathotype grouping of *S. endobioticum* isolates is based on tuber-based bioassays but agroinfiltration experiments are typically performed on leaves. We previously demonstrated that plant *R* genes for the EPPO differential set (6) were equally expressed in aboveground plant parts compared to the tuber-based assays (35). This justifies the use of the leaf-based agroinfiltration assay to test the interaction between candidate *Avr* genes and *R* genes.

### AvrSen1

As the interaction between *S. endobioticum* and its host is specific, the *AvrSen1* gene was hypothesized to be present in the species specific secretome. Indeed the identified *AvrSen1* gene had no orthologs in the closely relate species *S. microbalum* which has a non-pathogenic saprobic lifestyle.

Cartwright (1926) reported that in incompatible interactions, zoospores encyst on the host cell and penetration occurs. Initially the fungal thallus develops normally, but eventually an immune response is triggered resulting in localized cell death (18). This observation suggests that the strategy of overcoming pathogen/microbe associated molecular patterns (PAMP) triggered immunity (PTI) by *S. endobioticum* may be identical in the different pathotypes and that the differentiation between pathotypes is the result of effectors secreted in the cytoplasm after host penetration. The strong response of Sen1 plants to the cytoplasmic AvrSen1ΔSP construct, and absence of a response when the *AvrSen1* was expressed with signal peptide, supports the intracellular recognition of the effector. Also, no nuclear localization signals, chloroplast or mitochondrial targeting peptides were present, suggesting that the protein is not localized in these subcellular compartments, but has a cytoplasmic target recognition. Similar observations were made for the obligate biotrophic flax rust (*Melampsora lini*) effectors *AvrM* and *AvrL567* (36, 37). We hypothesize that also in case of wart resistance genes NLR immune receptors are involved. In flax it are cytoplasmic NLR immune receptors that recognize the cognate *M. lini Avrs*. Many of the described fungal *Avr* genes lack functional domains and have no predicted function (38). Similarly, no Pfam domains, GO-terms or protein family membership were predicted for AvrSen1, and its function in virulence remains elusive.

The *AvrSen1* gene (SeMB42_g04087 from the MB42 genome sequence) has a single ortholog in the Canadian pathotype 6(O1) isolate LEV6574 genome, i.e. SeLEV6574_g04683. Both genomes were independently sequenced, assembled and annotated (31), and the gene prediction in the LEV6574 genome reflects the truncated *AvrSen1* variant 2 that was identified from the *AvrSen1* prediction pipeline. This indicates that not only the particular SNP is present in the Canadian genome, but also that the truncated gene model is expressed (Fig. S6). Being a single copy ortholog in both *S. endobioticum* isolates is atypical for an *Avr* gene as many effectors belong to multi-gene families and have diversified from a common ancestor (20). In addition, the number of cysteine residues (2) is lower, and the protein length (376 amino acids) is larger compared to the features generally attributed to effector proteins. We would have not been able to detect the *AvrSen1* gene when applying these features as selection criteria as was suggested by others (32).

### *AvrSen1* and its variants in *S. endobioticum* isolates

Five variants of the *AvrSen1* gene were identified, which suggests that the *AvrSen1* gene is under strong Sen1 selection pressure that is mainly exerted in potato cultivation. Indeed Sen1 is widely deployed in current potato varieties (39). In contrast, 79% of all 8,031 *S. endobioticum* genes included in the screening for *AvrSen1* candidates do not display any dN changes or reduced gene coverage for any of the fourteen isolates included in this study. A similar observation was made by Huang *et al.* (40) who analyzed genetic variations of six *Avr* genes in the rice blast fungus *Magnaporthe oryzae*, and compared these to seven randomly selected non-*Avr* control genes. In *M. oryzae*, Avr genes frequently show deletions and high levels of nucleotide variation leading to (shared) non-synonymous substitution in the diversified rice blast strains. Of the five *AvrSen1* variants, three are truncated gene models as the result of single nucleotide insertions or substitutions. Similarly, in the oomycete *P. infestans* a truncated version of the Avr4 protein remains unrecognized by plants with the *R4* gene (41).

Variant 1 (avrSen1:Asp^256^>stop^256^) was found most frequently in higher pathotypes representing different mitochondrial haplogroups which are believed to be independently introduced in Europe from the Andes (10). Hence it is likely that the mutation leading to avrSen1:Asp^256^>stop^256^ is emerged early in the evolution and spread of the pathogen, which would suggest that the *Sen1* locus should also be present in (wild) potato varieties in the native range of *S. endobioticum*. Variant 2 (avrSen1:Gln^306^>stop^306^) was found exclusively in the four Canadian pathotype 6(O1) isolates. *S. endobioticum* was introduced from Europe to Canada in the early 1900s to Newfoundland (12) from which it spread to Saint Edwards Island from where the sequenced isolates were obtained. The variant 2 haplotype was not found in the European isolates, which could be an effect of sampling, or it could indicate that the SNP leading to variant 2 is recent and occurred in the Canadian *S. endobioticum* isolates *de novo*. In South America, the presumed center of origin of potato, *S. endobioticum* interacts with many other *Solanum* species and their respective resistance genes. In this respect it is interesting to note that the Peruvian isolate has retained the intact *AvrSen1*.

### Disruptive selection

After two multiplications of pathotype 1(D1) isolate 01WS on potato cultivar Erika, which does not provide full resistance to 1(D1), wart formation was obtained. The fungal isolate from the warted tissues (E/II/2015) was pathotyped, and produced a pathotype 6(O1) phenotype. This sample could be multiplied on (among others) Producent which contains *Sen1* (Prodhomme *et al.*, unpublished). From the amplification of the *AvrSen1* gene followed by PacBio SMRT sequencing we expected to observe a loss of the *AvrSen1* gene as a result of a selection for loss of function mutation induced by the partially resistance cultivar Erika. Indeed we observed a loss of *AvrSen1* in isolate E/II/2015, but to our surprise a novel variant was found: variant 5.

Variant 5 is the result of a substitution on the first base of the third intron in the *AvrSen1* gene sequence. The first two bases of the exon-intron boundary are highly conserved and are required for splicing of the introns out the pre-mRNA (42). Polymorphisms in this sequence results in a read-through of the third exon resulting in the introduction of a stop codon at the 94^th^ amino acid position. Low levels (0.0006%) of the SNP leading to *AvrSen1* variant 5 in isolate E/II/2015, were identified in isolate 01WS. However, at these low percentages it is impossible to differentiate between sequence errors introduced by PacBio SMRT sequencing, errors introduced by amplification errors, or a true genotype. Nonetheless, the CCS reads with the variant 5 haplotype in isolate 01WS were of high quality and were built from 4 to 125 passages with an average of 39 passages. Analysis of within-isolate diversity using mitochondrial haplotypes showed increased diversity in isolate E/II/2015 relative toto the original pathotype 1(D1) isolate. Selection against the main genotype allowed proliferation of individuals (10).

Where in isolate E/II/2015 almost all variation (99.3%) was selected against the *AvrSen1* allele, low frequencies of *AvrSen1* (7.0 to 8.2%) were identified in variant 1 isolates 02WS, MB56, SE4, SE5, and SE6. These variant 1 isolates were multiplied on the universal susceptible variety Deodara which does not contain *Sen1. AvrSen1* is found in most isolates tested, although sometimes in a minor haplotype. Both alleles could be maintained in the population on account of balancing selection, and have a positive effect on the overall fitness of the population.

The observed within-isolate variation, with low levels of the avirulent allele being present, could explain the escapes in susceptible pathogen-host interactions which have been reported in *S. endobioticum* bioassays (43). This is further supported by the observation that in a *S. endobioticum* bioassay both wart formation and a HR can be observed on different shoots of the same tuber cutting.

### Closing remarks and outlook

Here we report the identification of the first *Avr* gene in Chytridiomycota which demonstrates that the gene-for-gene model applies to the potato-*S. endobioticum* pathosystem. Our strategy proved to be effective in identifying the *AvrSen1* gene and a similar approach may result in uncovering more *Avr* genes. This is particularly true when it is specifically matched to QRL or *R* gene based predicted absences and presences. *AvrSen1* could also be instrumental in finding *Avr* genes in other chytrid pathogens such as *B. dendrobatidis*, and may also help to identify the potato *Sen1* gene.

The *AvrSen1* gene is under strong selective pressure and several forms of loss of the *AvrSen1* locus were observed. Given the broad presence of variant 1, and to a lesser extend variant 2, in non-pathotype 1(D1) isolates, these could potentially be recognized by other *R* genes, making them avirulence factors. Since most mutations resulted in a C-terminal truncation, this suggests that this part is essential for recognition by Sen1. Additional research is required to show which part of AvrSen1 is indeed recognized.

We previously observed that pathotypes 2(G1) and 6(O1) isolates that produced the same phenotype were present in different mitochondrial haplogroups (10), and concluded that these phenotypes could have emerged independently from different genetic backgrounds. Our results regarding the different types of loss of *AvrSen1* in the pathotype 2(G1) and 6(O1) isolates further strengthens this hypothesis (Fig. 4). Additionally, when regarding all species specific secreted proteins as potential effectors, different patterns of predicted loss of function can be seen between the two pathotype 2(G1) isolates, and the Canadian and European pathotype 6(O1) isolates. This analysis also allows further differentiation between the two pathotype 1(D1) isolates (Fig. S7). These different types of pathotype 2(G1) and 6(O1) could have evolved their own set of *Avr* genes which are not detected in the current pathotyping methods as the cognate *R* genes are not included in the differential potato panel. The *AvrSen1* gene can be used to screen isolates to identify these possible different genotypes in isolates phenotypically identified as pathotype 1(D1). In time, functional markers such as the *AvrSen1* can contribute to alternative pathotyping assays.

## Materials and methods

### Identification of *AvrSen1* candidates

Whole genome sequence data used in this study were generated by Van de Vossenberg *et al.* (10) and were obtained from resting spores extracted from fresh warts (table S1). These datasets comprised 30 isolates that were grouped into seven different pathotypes. Datasets with >10x median coverage and known pathotype identity were included in this study (Fig. S1). Sequence reads were mapped (length fraction: 0.8, similarity fraction: 0.9) to the *S. endobioticum* pathotype 1(D1) isolate MB42 reference genome (31) in CLC genomics workbench v11.0.1 (Qiagen, the Netherlands), and mappings were improved with the local realignment tool (default settings) to better resolve the mapping in areas around insertions and deletions.

To determine putative amino acid sequence polymorphisms in different isolates, variants were called using the basic variant detection tool (min_coverage_: 5, min_count_: 3, Min_frequency_: 70%, broken reads: ignore), and non-synonymous (dN) changes were identified with the Amino Acid Changes tool. Using this tool, the number of dN changes relative to the MB42 genome were determined per gene for each of the isolates tested. Genes with one or more dN change in an isolate were regarded to be putatively functionally absent (i.e. loss of function). To determine (partial) deletions of genes in different isolates, the percentage of length coverage relative to the MB42 reference genome was determined. Per mapped isolate, a consensus sequence was created for each gene, and the number of nucleotides with no coverage were determined. Genes with less than 90% coverage in an isolate were regarded to be putatively structurally absent. Genes present in pathotype 1(D1) isolates, but absent in higher pathotypes were regarded as candidate *AvrSen1* genes.

These candidate genes were individually inspected to verify if they were legitimate candidates by determining: a. correctness of gene prediction by checking the MB42 RNAseq (31) read mapping to the MB42 reference genome; b. significance of dN change and variants present in other pathotypes by differentiating between conservative and non-conservative dN changes; c. significance of <90% length coverage, and d. functional predictions with InterProScan. Presence of nuclear localization signals, chloroplast or mitochondrial targeting peptides in AvrSen1 were determined with LOCALIZER v1.0.4 (44). A graphical summary of detection pipeline is presented in Figure 1.

### The *S. endobioticum* specific secretome

The secretome was defined as proteins possessing a secretion signal as predicted with SignalP v4.1 (45), absence of a mitochondrial targeting peptide as determined with TargetP v1.1 (46) and absence of transmembrane helices and/or GPI anchors as determined with TMHMM v2.0 (47) and PredGPI (48) respectively. Next, OrthoFinder v1.1.4 (49) output generated by Van de Vossenberg & Warris *et al.* (31) was used to identify *S. endobioticum* specific genes. In short, protein sequences of the *S. endobioticum* pathotype 1(D1) MB42 and pathotype 6(O1) LEV6574 reference genomes were compared to the proteomes of nine other chytrid isolates, including the closely related saprobic *Synchytrium microbalum* and the more distant amphibian decline pathogen *Batrachochytrium dendrobatidis*. From this analysis, orthologous groups unique to both *S. endobioticum* isolates or isolate MB42 were regarded *S. endobioticum* or pathotype 1(D1) specific.

### Cloning of *AvrSen1* candidates and variants

Coding sequences of the *AvrSen1* candidate and its truncated variants were synthesized by GenScript (China) after codon optimization for plants and removal of *BsaI, BpiI*, and *BsmBI* restriction sites. Gene expression constructs were prepared with the Golden Gate Cloning system and contained the *AvrSen1*, avrSen1:Asp^256^>stop^256^, avrSen1:Gln^306^>stop^306^ coding sequences inserted between the Cauliflower mosaic virus (CaMV) plCSL13001 promotor + 5’UTR and the CaMV plCH41414 3’UTR + terminator sequences (50) in a modified pBINPLUS binary vector (pBINPLUS-GG). *AvrSen1* constructs were prepared with and without the *Nicotiana benthamiana* CRT (calreticulin) signal peptide plCH37326 to test for differential apoplastic or cytoplasmic recognition. The correctness of the constructs was verified using Sanger sequencing. pBINPLUS-GG plasmids with *AvrSen1* inserts were transformed to *A. tumefaciens* strain AGL + VirG (51) using electroporation. The presence of the intact plasmid was confirmed using PCR, before using the transformed colonies for agroinfiltration.

### Agroinfiltration

The verified *AvrSen1* candidate and two of the functional variants were cloned and transiently expressed in potato leaves to determine if their gene products were recognized by *Sen1*. Progeny from crosses between the pathotype 1(D1) resistant varieties Desiree and Kuras and the pathotype 1(D1) susceptible variety Aventra were selected (Prodhomme *et al.*, unpublished), based on their resistance or susceptibility to *S. endobioticum* pathotype 1(D1), on the presence or absence of *Sen1*, and on their competence for transient expression.

All progeny clones and the parent varieties were field propagated to provide tubers. In 2016, six and eight tubers respectively, were tested for pathotype 1(D1) resistance using the Glynne-Lemmerzahl and Spieckermann bioassays (52). A subset of clones and the parents were re-phenotyped in 2017 with 6 and 12 tubers respectively. Presence or absence of *Sen1* was determined using five KASP markers specific to the *Sen1* haplotype (Prodhomme *et al*., unpublished). The progeny clones were grown *in vitro* and tested for Agrocompetence with *A. tumefaciens* suspensions at OD 0.3 and 0.1. A positive control consisting of the co-infiltration of the *P. infestans Avr8* gene and the cognate *R8* gene [38]) and a negative control (GUS) were used in this experiment. Non-competent lines which showed aspecific reactions to *A. tumefaciens* or did not show any HR when infiltrated with the positive control were excluded.

The transformed colonies were cultured in 5 ml of LB medium containing the appropriate antibiotics and grown overnight (28 - 30 °C, shaking 200 rpm). Between 20 to 200 μL of the LB cultures was diluted in 15 mL YEB medium containing the appropriate antibiotics, 1.5 μL acetosyringone (200 mM) and 150 μL MES (2-(N-morpholino)-ethane sulfonic acid, 1 M) and grown overnight (28 - 30 °C, shaking 200 rpm). The YEB cultures were centrifuged for 10 minutes at 4000 x g. The supernatant was poured off and the pellet carefully re-suspended until the appropriate OD in freshly made MMA containing 1 ml/L of acetosyringone. The cultures were incubated 1 to 3 hours before infiltrations at room temperature in the dark.

Potato plants were clonally propagated from *in vitro* culture on Murashige & Skoogs medium supplemented with 2% Sucrose. Two weeks after incubation at 25 °C, the rooted cutlings were transferred to the greenhouse in 11 cm pots with potting soil under greenhouse conditions (constant 18 to 21 °C and 90% relative humidity with a light regime of 16 h light /8 h darkness). Using a syringe, small amounts of bacterial suspensions (OD 0.3 or 0.1) containing the pBINPLUS-GG with inserts of interest, were injected on the abaxial side of selected leaves of 3 to 4 weeks after planting in the greenhouse. Each genotype was tested in duplicate and per plant three leaves were injected, referred to as “low”, “middle”, and “high” based on their relative position on the plant. On each leaf, a positive control (*Avr8*/*R8* co-infiltration (53)) and a negative control (GUS or *R8* gene) was included to monitor the effectiveness of the Agroinfiltration experiments. After injection, plants were kept in the greenhouse for 2-5 days before scoring HR reactions following Rietman *et al*. (54).

### *AvrSen1* and its variants in *S. endobioticum* isolates

The presence of functional *AvrSen1* genes in sequenced isolates was superimposed to the *S. endobioticum* haplotype network as described by Van de Vossenberg *et al*. (10), which was based on complete mitochondrial genomes. Isolates in the network were colored according to presence of *AvrSen1* or its functional variants, which was based on the Illumina read-mappings of 30 isolates (identification of AvrSen1 candidates) and PacBio SMRT sequencing of AvrSen1 amplicons (see below).

### Co-occurrence of *AvrSen1* and its variants within *S. endobioticum* isolates

As *S. endobioticum* isolates may contain a population of genotypes (10), and we observed allelic variation for *AvrSen1* in the Illumina sequences derived from a single isolate, we assessed the variation of *AvrSen1* within isolates by next generation sequencing of *AvrSen1* amplicons. Isolates used by Van de Vossenberg *et al.* (10) representing different pathotypes and mitochondrial haplogroups were selected for the analysis, including the German pathotype 1(D1) isolate 01WS and laboratory isolate E/II/2015 (table S2). The latter isolate was produced after two multiplications of 01WS on the semi-resistant variety Erika after which it showed a pathotype 6(O1)-like phenotype.

Primers were designed allowing generic amplification of the *AvrSen1* locus (Fig. S2). Amplification of *AvrSen1* gene was performed in 20 μL reaction mixes based on the Takara Premix HotStart (TaKaRa, Japan) reagents containing 500 nM of a tagged AvrSen1_fw (*tag*-5’-CTG GAA GCT CTA TTT CAT AGG TCA-3’) primer, 500 nM of a tagged AvrSen1_rv (*tag*-5’-CAC TCA CTC GTG CCA TTT CTA-3’) primer and 2 μL target DNA. PCR thermocycler conditions were as follows: 2 min. 94 °C initial denaturation followed by 35 cycles of (30 s. 94 °C, 30 s. 55 °C, and 2 min. 72 °C) and with a final elongation of 5 min. at 72 °C. Amplification primers were tagged to create sample specific combinations allowing selection of isolate specific sequence data on the pooled amplicons. Amplicons were pooled in equimolar amounts, and subjected to PacBio SMRT sequencing. From the raw PacBio sequence data, circular consensus sequences (CCS) were generated and successively, CCS data were binned based on their sample specific tags using a custom script (https://git.wur.nl/brank001/pacbio-demultiplexing). Sample specific CCS data were mapped to the *AvrSen1* gene in Geneious Prime (55) (Biomatters Limited, New Zealand) using a reference assembly with default settings. Next, variants were detected and quantified with the Find Variations/SNPs tool (Minimum variant frequency = 0.1, Maximum variant P-value = 10^-4^).

## Acknowledgements

We thank Averis seeds BV, Böhm-Nordkartoffel Agrarproduktion GmbH & Co. OHG, Danespo, HZPC Holland BV, C. Meijer BV, SaKa Pflanzenzucht GmbH & Co. KG and Teagasc for funding, biological material and stimulating scientific discussions.

## List of supplementary material

Figure S1. Median read coverage in thirty *S. endobioticum* Illumina read sets

Figure S2. Plasmid map

Figure S3. Design of amplification primers flanking AvrSen1 and its variants

Figure S4. Number of passages in PacBio SMRT CCS sequences generated from AvrSen1 amplicons

Figure S5. Detail of PacBio CCS reads generated for isolates 01WS and E/II/2015

Figure S6. RNAseq for the AvrSen1 gene in the Dutch pathotype 1(D1) isolate MB42 and truncated variant 2 in the Canadian pathotype 6(O1) isolate LEV6574

Figure S7. Clustering of the species specific secretome of twelve *S. endobioticum* isolates

Table S1. *Synchytrium endobioticum* isolates and NextGen sequence information

Table S2. Samples included in within-isolate variation analysis using PacBio sequencing of *AvrSen1* amplicons

Table S3. Secretome analysis statistics

Table S4. Statistics of putative functional and structural absence for 14 *S. endobioticum* datasets relative to the pathotype 1(D1) MB42 reference genome

Table S5. Progeny and parents of Aventra x Desiree and Aventra x Kuras crosses selected for agroinfiltration

## Author Contributions

BvdV, TvdL, JV produced the research outline and wrote the manuscript. BvdV, GvA, MvG, MB, and CP performed laboratory and greenhouse experiments. RV provided critical comments to the manuscript and arranged critical research infrastructure. BB provided bioinformatics support. JP provided essential research material.

